# Proteome profiling of cutaneous leishmaniasis lesions due to dermotropic *Leishmania donovani* in Sri Lanka

**DOI:** 10.1101/2024.01.07.574579

**Authors:** Nuwani H. Manamperi, Nimesha Madhushani Edirisinghe, Harshima Wijesinghe, Lakmali Pathiraja, Nishantha Pathirana, Vishmi Samudika Wanasinghe, Chamalka Gimhani de Silva, W. Abeyewickreme, Nadira D. Karunaweera

## Abstract

Characterization of the host response in cutaneous leishmaniasis (CL) through proteome profiling has gained limited insights in leishmaniasis research, in comparison to that of the parasite. The primary objective of this study was to comprehensively analyze the proteomic profile of the skin lesions tissues in patients with CL, by mass spectrometry, and subsequent validation of these findings through immunohistochemical methods. Sixty-seven proteins exhibited significant differential expression between tissues of CL lesions and healthy controls (p<0.01), representing numerous enriched biological processes within the lesion tissue, as evident by both the Kyoto Encyclopedia of Genes and Genomes (KEGG) and Reactome databases. Among these, the integrated endoplasmic reticulum stress response (IERSR) emerges as a pathway characterized by the up-regulated proteins in CL tissues compared to healthy skin. Expression of endoplasmic reticulum (ER) stress sensors, inositol-requiring enzyme-1 (IRE1), protein kinase RNA-like ER kinase (PERK), and activating transcription factor 6 (ATF6) in lesion tissue was validated by immunohistochemistry. In conclusion, proteomic profiling of skin lesions carried out as a discovery phase study revealed a multitude of probable immunological and pathological mechanisms operating in patients with CL in Sri Lanka, which needs to be further elaborated using more in-depth and targeted investigations.

**Author Summary:** Cutaneous leishmaniasis (CL), is a skin infection caused by a type of single-celled parasite. These parasites are usually transmitted through the bite of infected sandflies. In Sri Lanka, CL is caused by a parasite type that usually causes a more severe disease form, known as visceral leishmaniasis. Interaction between the parasite and the human host is important in determining the disease outcome and hence, we conducted a study to look at the proteins in the skin lesions of people with CL using a technique called mass spectrometry. We found 67 proteins that were different between CL lesions and healthy skin. These proteins are involved in various processes in the body, and one specific process called the integrated endoplasmic reticulum stress response (IERSR) was more active in CL patients. We confirmed this by studying specific proteins related to stress in the lesion tissue. In conclusion, our study uncovered several potential immune and disease-related mechanisms in CL patients in Sri Lanka. However, more detailed investigations are needed to fully understand these processes.

## Introduction

Human leishmaniasis, encompassing its principal clinical manifestations including cutaneous (CL), mucocutaneous (MCL), and visceral leishmaniasis (VL), poses a significant burden, particularly in tropical regions [1,2]. The intracellular protozoan parasite *Leishmania* serves as the etiological agent of leishmaniasis and the genus *Leishmania* contains approximately 21 species that can result in a range of clinical presentations in humans depending on the infecting species [3]. Over the last three decades, Sri Lanka has witnessed the emergence of CL as a parasitic disease caused by *Leishmania donovani* zymodeme MON-37 [4].

The term “Proteome” is defined as the entire set of proteins and their alternative forms in a specific species and “proteomics” is defined as a large-scale and comprehensive study of a certain proteome [5]. In the field of leishmaniasis, proteomic profiling has predominantly focused on analyzing the proteome of the parasite, while comprehensive protein profiling of the host is less commonly performed [6]. Proteomic profiling, coupled with genome annotation techniques, has led to the identification of novel genes associated with the virulence of *Leishmania major.* These findings underscore the importance of studying both the parasite and host proteomes to gain comprehensive insights into the pathogenesis and virulence mechanisms of leishmaniasis [7].

As gene expression regulation primarily occurs at the post-transcriptional level, it necessitates the use of proteomics to effectively identify stage-specific proteins. The identification of such proteins is instrumental in elucidating the dynamic changes occurring at each different stage of the parasite’s life cycle. Additionally, it is proposed that metabolomics, the study of small molecules involved in cellular metabolism, holds great promise in enhancing our comprehension of parasite biology, identifying key drug targets, and unraveling mechanisms of drug resistance. The integration of proteomic and metabolomic approaches will improve knowledge of the *Leishmania* parasite and will help the development of more effective therapeutic strategies [8]. According to Kumar *et al* proteomic profiling of *L. donovani* soluble proteins have identified several novel and hypothetical proteins which can be explored as new drug targets or vaccine candidates in VL [9].

Furthermore, a study done by Hajjaran *et al*. on *Leishmania tropica* has identified that most responsive proteins in the visceral isolate exhibited lower abundance in the cutaneous isolate. In this study the largest clusters comprised proteins associated with carbohydrate metabolism and protein synthesis. Notably, a significant proportion of the identified proteins implicated in energy metabolism, cell signaling, and virulence demonstrated down-regulation, whereas certain proteins involved in protein folding, antioxidant defense, and proteolysis exhibited up-regulation in the visceral form [3].

Proteomic profiling of cutaneous lesions due to *Leishmania* in humans remains limited in the available literature. However, a noteworthy study on proteome profiling in CL associated with *Leishmania braziliensis* revealed the up-regulation of caspase 9, along with the presence of caspase-3 and granzyme B in the lesions, which are known to contribute to the progression of tissue damage. Moreover, several biological functions, including apoptosis, immune response, and biosynthetic processes, were observed in both the lesions and healthy skin, but they were notably up-regulated in the lesions. The analysis of protein-protein interactions highlighted the cytotoxic T lymphocyte-mediated apoptosis of target cells as the main canonical pathway represented [6]. These findings shed light on the molecular mechanisms underlying CL lesions and provide valuable insights into the processes associated with tissue damage and immune responses in the context of *Leishmania* infections. Serum proteomic analysis in VL due to *L. donovani* has revealed differentially expressed serum proteins, which may be used as biomarkers of disease prognosis [10].

The current study aims to comprehensively analyze the proteomic profile of human CL lesions due to *Leishmania donovani* infections in Sri Lanka and validate the biological pathways represented.

## Results

### Proteomic profile in cutaneous leishmaniasis host tissues

A total of 1290 proteins were identified and after filtering those with a very low level of expression among the groups, 388 proteins were selected for analysis. After adjusting for multiple corrections, a total of 67 proteins (Table 1) were seen as differentially expressed between CL lesions and healthy controls (p<0.01). Among these proteins three proteins were down-regulated and 64 proteins were up-regulated.

**Table 1:** Proteins differentially expressed between cutaneous leishmaniasis lesions and normal skin.

| No | Gene name | Protein name | UniProt entry no. | Expression change | P value |
| --- | --- | --- | --- | --- | --- |
| 1. | OGN | Osteoglycin | P20774 | Down-regulated | 0.00004 |
| 2. | PSME1 | Proteasome activator subunit 1 | Q06323 | Up-regulated | 0.00006 |
| 3. | MYH9 | Myosin heavy chain 9 | P35579 | Up-regulated | 0.00016 |
| 4. | TMPO | Thymopoietin | P42167 | Up-regulated | 0.00017 |
| 5. | WARS | Tryptophanyl-tRNA synthetase | P23381 | Up-regulated | 0.00018 |
| 6. | HMGB2 | High mobility group protein B2 | P26583 | Up-regulated | 0.00026 |
| 7. | PDIA3 | Protein disulfide isomerase family A, member 3 | P30101 | Up-regulated | 0.00026 |
| 8. | PRKCSH | Glucosidase II subunit beta | P14314 | Up-regulated | 0.00026 |
| 9. | HSPE1 | Heat shock 10kDa protein 1 | P61604 | Up-regulated | 0.00032 |
| 10. | COTL1 | Coactosin-like protein1 | Q14019 | Up-regulated | 0.00034 |
| 11. | LAP3 | Leucine aminopeptidase 3 | P28838 | Up-regulated | 0.00034 |
| 12. | RPL6 | Ribosomal protein L6 | Q02878 | Up-regulated | 0.00034 |
| 13. | MSN | Moesin | P26038 | Up-regulated | 0.00034 |
| 14. | IVL | Involucrin | P07476 | Up regulated | 0.00053 |
| 15. | TYMP | Thymidine phosphorylase | P19971 | Up-regulated | 0.00053 |
| 16. | CALM1 | Calmodulin1 | P0DP23 | Up-regulated | 0.00068 |
| 17. | HMGB1 | High mobility group protein B1 | P09429 | Up-regulated | 0.00068 |
| 18. | S100A9 | S100 calcium-binding protein A9 | P06702 | Up-regulated | 0.00078 |
| 19. | MZB1 | Marginal zone B and B1 cell-specific protein | Q8WU39 | Up-regulated | 0.00087 |
| 20. | CORO1A | Coronin, actin-binding protein, 1A | P31146 | Up-regulated | 0.00131 |
| 21. | TNC | Tenascin C | P24821 | Up- regulated | 0.00131 |
| 22. | KRT17 | Keratin 17 | Q04695 | Up-regulated | 0.00140 |
| 23. | HNRNPM | Heterogeneous nuclear ribonucleoprotein<br>M | P52272 | Up-regulated | 0.00160 |
| 24. | ARHGDIB | Rho GDP<br>dissociation<br>inhibitor (GDI)<br>beta | P52566 | Up-regulated | 0.00163 |
| 25. | RPL8 | Ribosomal protein<br>L8 | P62917 | Up-regulated | 0.00163 |
| 26. | SSB | Sjogren syndrome<br>antigen B (Lupas<br>La protein) | P05455 | Up-regulated | 0.00163 |
| 27. | KRT6A | Keratin 6A | P02538 | Up-regulated | 0.00163 |
| 28. | CANX | Calnexin | P27824 | Up-regulated | 0.002 |
| 29. | RPL4 | Ribosomal protein<br>L4 | P36578 | Up-regulated | 0.002 |
| 30. | TPM4 | Tropomyosin 4 | P67936 | Up-regulated | 0.0021 |
| 31. | HSPA5 | Heat shock 70kDa<br>protein 5 | P11021 | Up-regulated | 0.0022 |
| 32. | ICAM1 | Intercellular<br>adhesion<br>molecule 1 | P05362 | Up-regulated | 0.0023 |
| 33. | KRT6C | Keratin 6C | P48668 | Up-regulated | 0.0030 |
| 34. | HCLS1 | Hematopoietic<br>lineage cell-specific protein | P14317 | Up-regulated | 0.0031 |
| 35. | RPL35 | Ribosomal protein<br>L35 | P42766 | Up-regulated | 0.0033 |
| 36. | ALYREF | Aly/REF export<br>factor | Q86V81 | Up-regulated | 0.0034 |
| 37. | ALDOA | Aldolase A | P04075 | Up-regulated | 0.0037 |
| 38. | EEF2 | Eukaryotic translation elongation factor 2 | P13639 | Down-regulated | 0.0037 |
| 39. | FLG2 | Filaggrin family member 2 | Q5D862 | Up-regulated | 0.0038 |
| 40. | RPL13 | Ribosomal protein L13 | P26373 | Up-regulated | 0.0044 |
| 41. | NAP1L1 | Nucleosome assembly protein 1-like 1 | P55209 | Up-regulated | 0.0045 |
| 42. | LCP1 | Lymphocyte cytosolic protein 1 | P13796 | Up-regulated | 0.0049 |
| 43. | THRAP3 | Thyroid hormone receptor-associated protein 3 | Q9W2Y1 | Up-regulated | 0.0049 |
| 44. | TPM3 | Tropomyosin 3 | P06753 | Up-regulated | 0.0049 |
| 45. | PDIA6 | Protein disulfide isomerase family A, member 6 | Q15084 | Up-regulated | 0.005 |
| 46. | GBP1 | Guanylate binding protein 1 | P32455 | Up-regulated | 0.005 |
| 47. | LSP1 | Lymphocyte-specific protein 1 | P33241 | Up-regulated | 0.005 |
| 48. | ERP29 | Endoplasmic reticulum protein 29 | P30040 | Up-regulated | 0.0063 |
| 49. | STAT1 | Signal transducer and activator of transcription 1 | P42224 | Up-regulated | 0.0063 |
| 50. | TTR | Transthyretin | P02766 | Down-regulated | 0.0063 |
| 51. | CALR | Calreticulin | P27797 | Up-regulated | 0.0064 |
| 52. | NCL | Nucleolin | P19338 | Up-regulated | 0.007 |
| 53. | FBP1 | Fructose-1,6-bisphosphatase 1 | P09467 | Up-regulated | 0.0082 |
| 54. | AP3D1 | Adaptor-related protein complex 3, delta 1 subunit | O14617 | Up-regulated | 0.0082 |
| 55. | HNRNPC | Heterogeneous nuclear ribonucleoprotein C (C1/C2) | P07910 | Up-regulated | 0.0082 |
| 56. | LMNB1 | Lamin B1 | P20700 | Up-regulated | 0.0082 |
| 57. | HLA-A | Major histocompatibility complex, class I, A | P04439 | Upregulated | 0.0082 |
| 58. | P4HB | Prolyl 4-hydroxylase, beta polypeptide | P07237 | Up-regulated | 0.0084 |
| 59. | RPL5 | Ribosomal protein L5 | P46777 | Up-regulated | 0.0086 |
| 60. | RRBP1 | Ribosome binding protein 1 | Q9P2E9 | Up-regulated | 0.0086 |
| 61. | HNRNPU | Heterogeneous nuclear ribonucleoprotein U | Q00839 | Up-regulated | 0.009 |
| 62. | SFPQ | Splicing factor<br>proline/glutamine-<br>rich | P23246 | Up regulated | 0.009 |
| 63. | RPS16 | Ribosomal protein<br>S16 | P62249 | Up-regulated | 0.0092 |
| 64. | RPS19 | Ribosomal protein<br>S19 | P39019 | Up-regulated | 0.0092 |
| 65. | S100A4 | S100 calcium-binding protein<br>A4 | P26447 | Up-regulated | 0.0092 |
| 66. | FTL | Ferritin, a light<br>polypeptide | P02792 | Up-regulated | 0.00923 |
| 67. | KRT16 | Keratin 16 | P08779 | Up-regulated | 0.0094 |

### Protein–protein interaction analysis

Protein-protein interaction network for the differentially expressed proteins between CL lesions and healthy skin demonstrated 67 nodes and 116 edges with a protein-protein interaction p value of 1×10^−16^ which showed in Fig 1.

**Fig 1.**
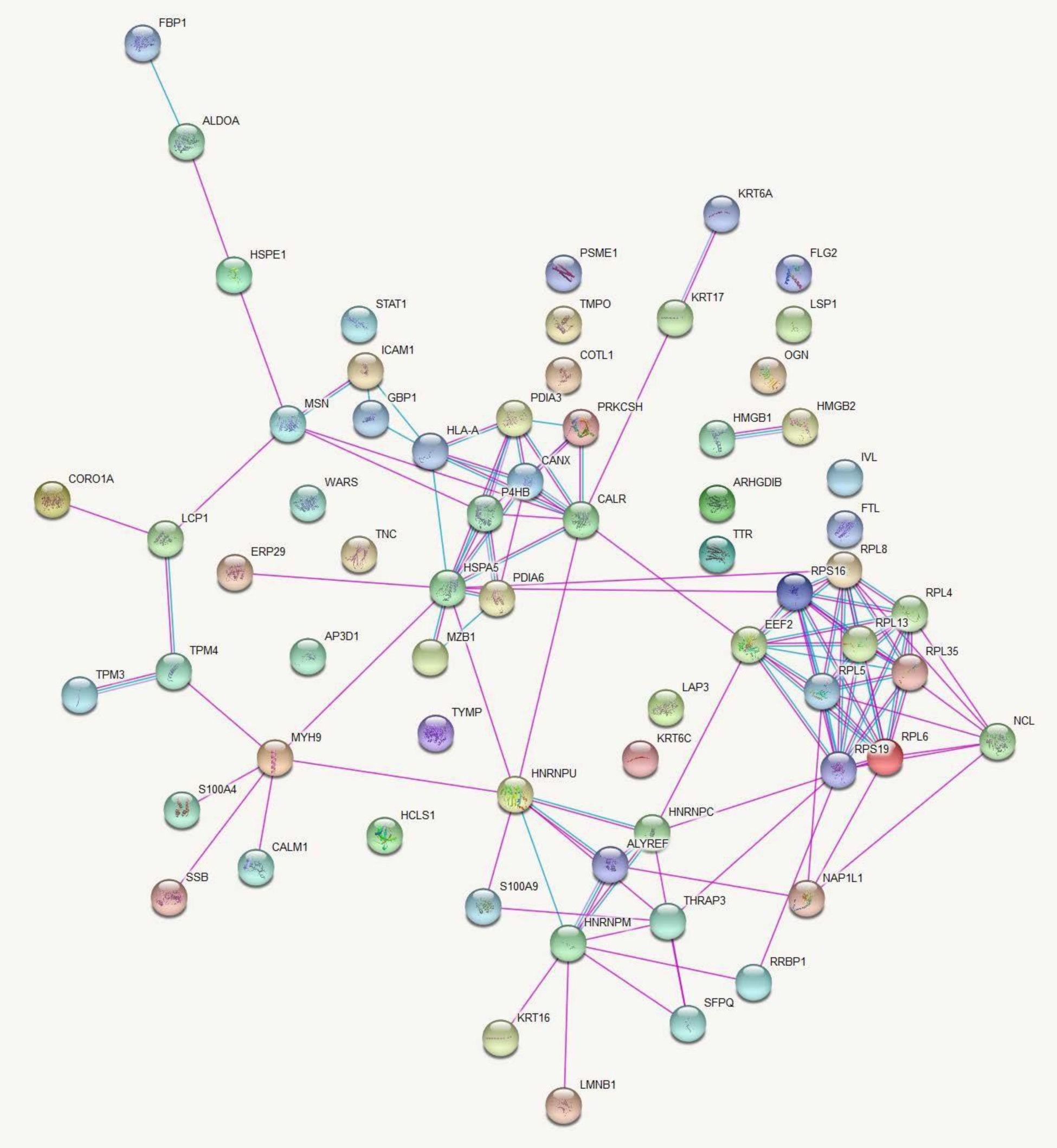
Networks of protein-protein interactions between differentially expressed proteins in CL patients and healthy controls. Proteins are represented by nodes and protein-protein interactions by edges (curated in pink and experimentally determined in blue). Filled nodes indicate that some 3D structure is known or predicted, whereas empty nodes represent proteins of unknown 3D structure. Figure credit: STRING.

Data analysis in this database provided a list of enriched pathways categorized based on Gene Ontology (GO) terms and Kyoto Encyclopedia of Genes and Genomes (KEGG) pathways (S1-S4 Tables). Protein-protein interactions in the KEGG pathways are significantly represented are illustrated in Figs 2 and 3.

**Fig 2.**
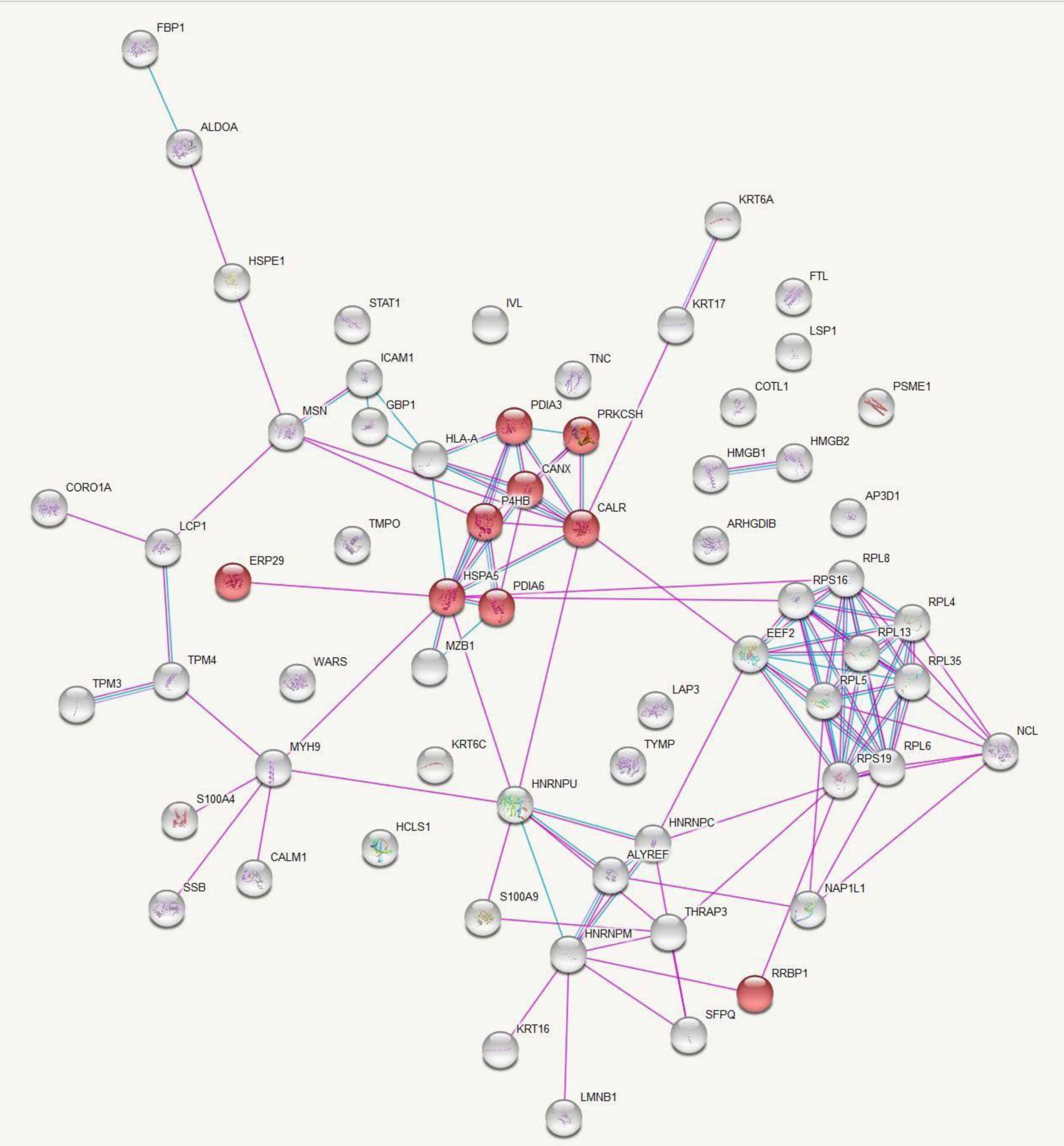
KEGG Pathway Analysis of Up-Regulated Protein Interactions in Leishmaniasis: Endoplasmic Reticulum Processing Insights. Protein-protein interactions among the proteins significantly up-regulated in leishmaniasis patients compared to healthy controls with proteins involved in the KEGG pathway ‘Protein processing in endoplasmic reticulum’ represented in red. Figure credit: STRING.

**Fig 3.**
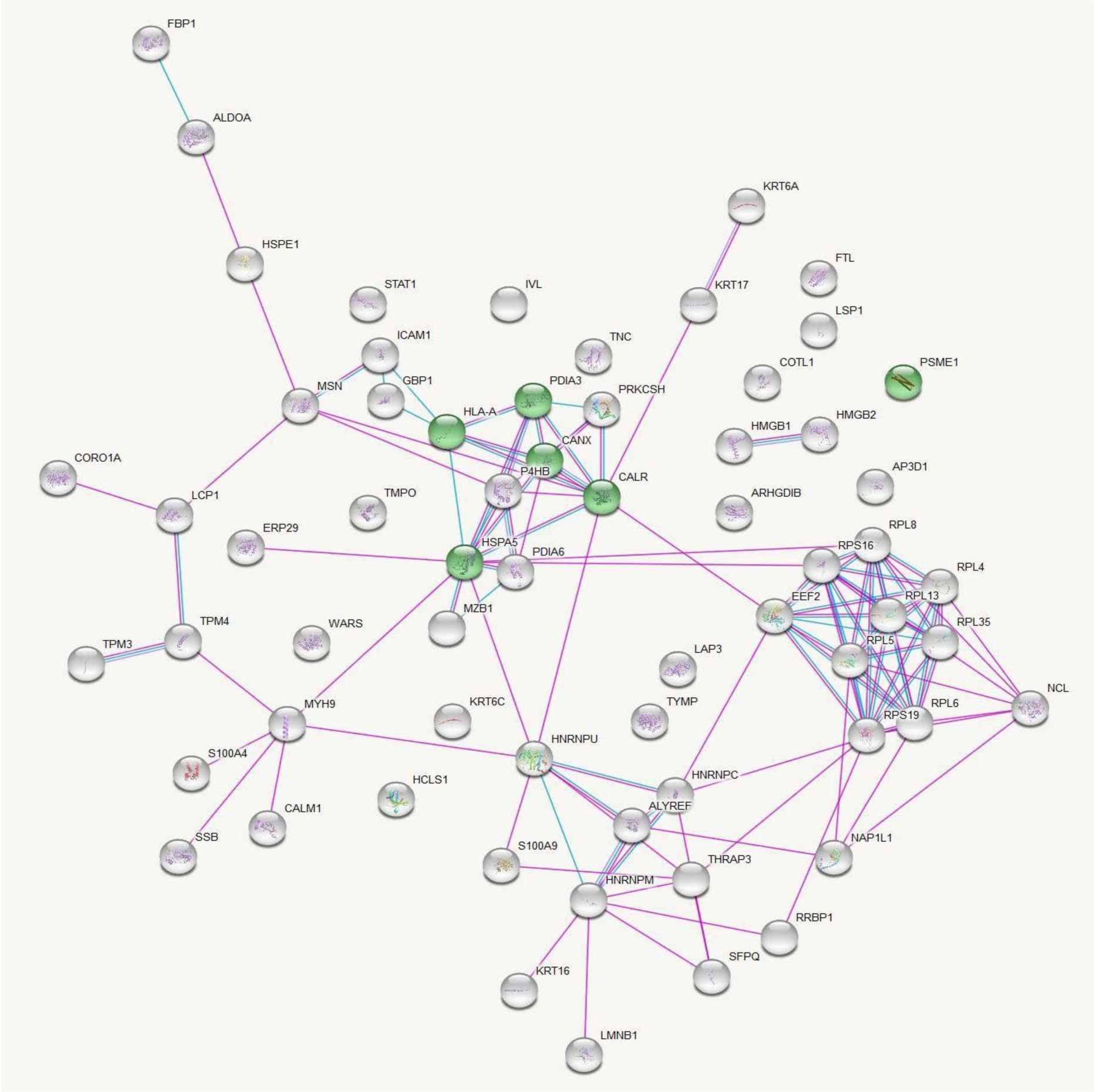
KEGG Pathway Analysis of Up-Regulated Protein Interactions in Leishmaniasis: Antigen processing and presentation insights. Protein-protein interactions among the proteins significantly up regulated in leishmaniasis patients compared to healthy controls with proteins involved in the KEGG pathway ‘Antigen processing and presentation’ represented in green. Figure credit: STRING.

### Pathway analysis for differentially expressed proteins by Reactome database

Gene names of significantly up regulated proteins were submitted to the Reactome pathway portal, version 3.2 (http://www.reactome.org) to identify pathways associated with these proteins. Pathways with ‘entities p-value’ <0.01 were considered as significantly associated with cutaneous leishmaniasis lesions (S5 Table). Submitted entities are as described in Table 1.

### Immunohistochemical validation of endoplasmic reticulum stress response

Most cases showed evidence of endoplasmic reticulum stress response on immunohistochemistry. Representative images in Fig 4 illustrated the staining patterns observed. Out of thirty cases examined, 13 (43.33%) cases showed positivity for only one of the three markers assessed (IRE1, PERK, ATF-6) and 17 cases (56.67%) showed positivity for all three. The details of the expression of the individual markers are shown in Table 2. Chi-square analysis revealed a statistically significant association (p<0.05) between the expression levels of IRE1, PERK, and ATF6 markers and both gender and histological grading in CL tissue samples. Histological grading was categorized as follows: 1) diffuse inflammatory infiltrate with parasitized macrophages, lymphocytes, and plasma cells, 2) parasitized macrophages with lymphocytes, plasma cells, and ill-formed histiocytic granulomata, and 3) a mixture of macrophages (with or without parasites), lymphocytes, plasma cells, and epithelioid granulomata.

**Fig 4:**
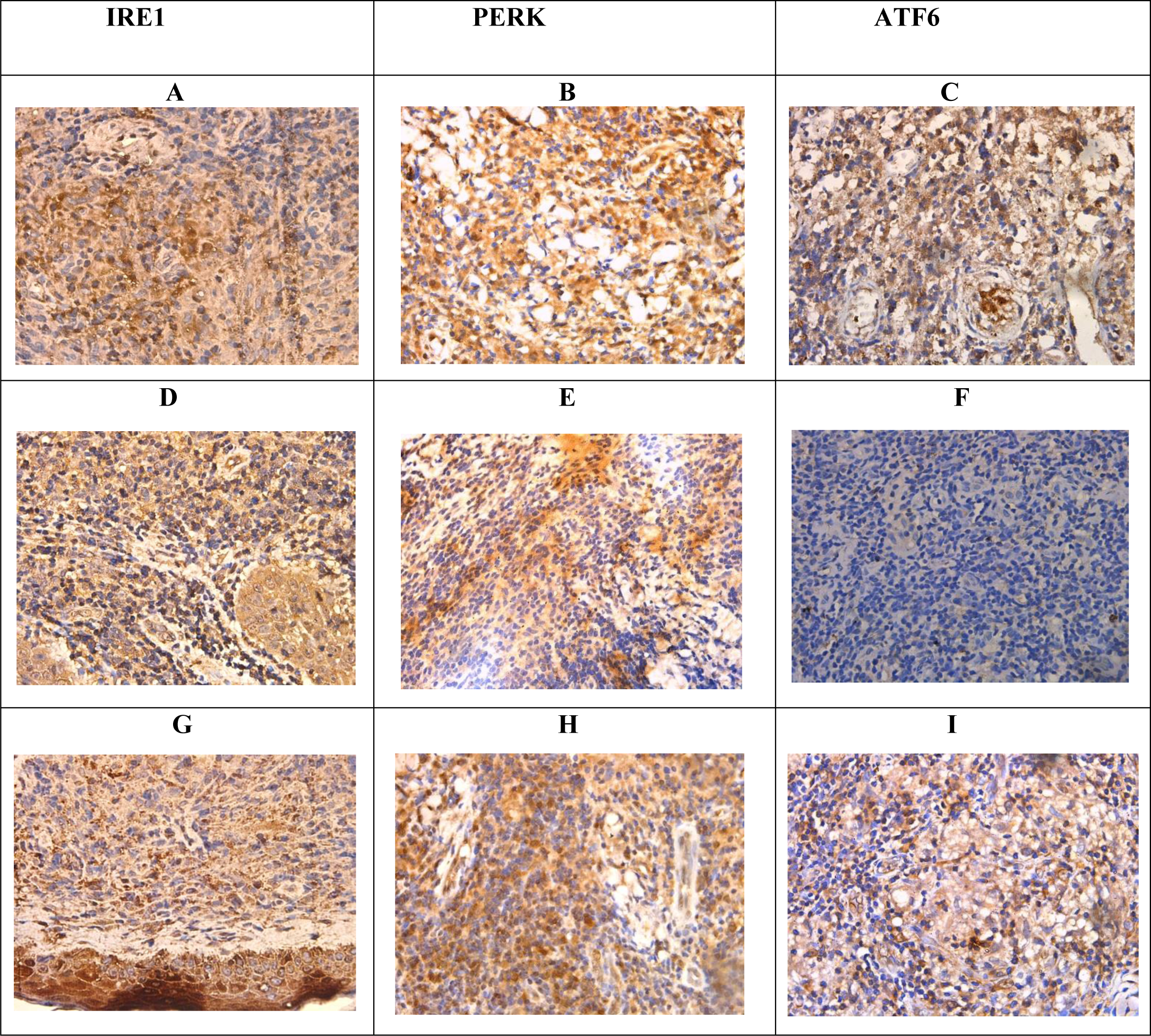
Immunohistochemistry for IRE1, PERK, and ATF6 in samples and controls. (A). IRE1-patchy nucleus and cytoplasmic positive (D), (G) nucleus negative and cytoplasmic positive. (B). PERK-focal nucleus and diffuse cytoplasmic positive. (E) nucleus negative and cytoplasmic positive (H) patchy nucleus and cytoplasmic positive. (C) ATF6 – nucleus negative and cytoplasmic positive (F) nucleus and cytoplasmic negative (I) focal cytoplasmic positive and nucleus (×400).

**Table 2:** Overall score for each marker of the study sample.

| ER Stress Markers | Overall Score |  |  |
| --- | --- | --- | --- |
|  | Negative | Low positivity | High positivity |
| IRE 1 | 8 (26.7%) | 15 (50%) | 7 (23.3%) |
| PERK | 2 (6.7%) | 15 (50%) | 13 (43.3%) |
| ATF6 | 7 (23.3%) | 10 (33.3%) | 13 (43.3%) |

## Discussion

Evaluation of parasite, host, and vector-associated factors are central to identifying the disease pathogenesis of CL in Sri Lanka, where cutaneous manifestations are the predominantly frequent disease presentation of a usually visceralizing parasite species. Proteomics profile data obtained from this study provides insights into disease pathogenesis, as shown by enriched biological processes and pathways related to local responses.

The integrated endoplasmic reticulum stress response (IERSR) is one such pathway represented by the up-regulated proteins in CL patients compared to healthy skin, determined by both the KEGG and Reactome databases. Protein processing in the endoplasmic reticulum (ER) includes glycosylation and folding of newly synthesized proteins with the help of luminal chaperons and transportation of correctly folded proteins in vesicles to the Golgi apparatus. Proteins that have not achieved proper folding are retained in the lumen of the endoplasmic reticulum, often in complex with molecular chaperones. When the protein synthesis and folding mechanism of the ER gets overwhelmed, misfolded proteins accumulate in the ER lumen, leading to a state of ER stress, which in turn activates a signaling pathway known as the unfolded protein response (UPR) to alleviate the ER stress. The three main stress sensors in the UPR: inositol-requiring enzyme-1 (IRE1) pathway, protein kinase RNA-like ER kinase (PERK) pathway, and activating transcription factor 6 (ATF6) pathway, induce downstream signaling cascades that make up the UPR. Although the UPR promotes cellular adaptations, if the ER stress is chronic these pathways signal towards cellular apoptosis. The IERSR or the UPR is proven to be associated with the pathology of diseases such as diabetes mellitus, neurodegeneration, inflammatory disorders, viral infection, and cancer [11]. Pathogens including bacteria and protozoa have the capacity to alter specific branches of UPR to avoid its detrimental effects.

Recent evidence suggests the involvement of specific arms of the UPR in disease pathogenesis in leishmaniasis due to several *Leishmania* species [12,13]. Emerging evidence suggests an important role for integrated ER stress response (IERSR) in the pathogenesis of *L. amazonensis* and *L. braziliensis* induced CL [14,15]. In *L. amazonensis* infection, induction of the IRE1/XBP1 arm was seen as beneficial for parasite survival by creating an environment with less oxidative stress, which is mediated by the increased IFN-β production. The PERK pathway is known to play a key role in autophagy and autophagy is suggested as a probable mechanism of supplying nutrition to the parasite and *L. amazonensis* is also known to induce autophagy in macrophages. It is also known that induction of a low level of stress may trigger an adaptive UPR, which increases the cellular resistance to subsequent ER stress, a process known as ER hormesis and IRE1 and PERK arms contribute to ER hormesis. Studies have shown that *L. infantum* is capable of inducing such a mild UPR in infected macrophages. Proteome profiling of CL in the present study has demonstrated a significantly increased expression of proteins associated with IRE1 and ATF6 pathways suggestive of an important role played by the UPR in the pathogenesis of *L. donovani* induced CL in Sri Lanka, which is further validated by immunohistochemical staining of the ER stress sensors in lesion tissues. In this study, significant difference was observed between the expression levels of IRE1, PERK, and ATF6 markers concerning both gender and histological grading. However, the exact role played by specific branches of the UPR in *L. donovani* infections needs to be further investigated.

Presence of an endobiont double-stranded RNA virus belonging to the *Totiviridae* family known as *Leishmaniavirus* (LRV) has been described in association with the *Leishmania (Viannia)* subgroup. The occurrence of a similar virus was described in a single isolate of *L. major* [16]. The presence of this virus has been associated with an increase in disease severity, parasite persistence, metastasis, and treatment failure [17]. Double-stranded RNA viruses are known to activate innate immune response via the Toll-like receptor (TLR)3 pathway which induces the secretion of IFN-β, which favors parasite survival [16]. To our knowledge, such an endobiont virus has not been demonstrated in *L. donovani*. Several pathways associated with viral infections such as viral gene expression and assembly of viral components at the budding site are represented by the upregulated proteins in CL patient samples in the present study, indicating the probability of having such an endobiont virus in the *L. donovani,* which should be further evaluated.

Furthermore, several pathways in the immune response such as class I MHC-mediated antigen processing and presentation, antigen processing with the cross presentation, interferon gamma signaling, IFN-α/β signaling, IL-12 family signaling, IL-6 signaling, IL-35 signaling, and neutrophil degranulation, leucocyte migration are represented by the significantly up-regulated proteins in patient samples. Antigen presentation by class I MHC is mainly restricted to proteins synthesized within the cell and hence play a major role in viral antigen presentation to CD8+ T cells. In the process of antigen cross-presentation, exogenous antigens are presented to CD8+ T cells in association with class I MHC molecules instead of with MHC class II molecules [18]. This is well known for infections with intracellular pathogens with the antigen-presenting cells (APC) becoming the major source of antigen cross-presentation. Antigen-presenting cells acquire these exogenous antigens from phagosomes (cell-associated antigens) or endosomes (soluble protein antigens). Antigenic proteins are processed and loaded onto class I MHC molecules. There are two pathways depending on the requirement for cytosolic proteases and a transporter associated with antigen processing (TAP). The cytosolic pathway is dependent on both TAP and proteasome, whereas the vacuolar pathway is independent of both. Processed peptides are loaded onto class I MHC molecules in the ER or the phagosome [19]. In *Leishmania* infections antigens are processed in phagosomes and cross-presented via class I MHC in a TAP-independent pathway to induce a cytotoxic T-cell response. In addition to this, antigens of *Leishmania* spp. are also presented to CD4+ T helper cells in association with class II MHC molecules [20]. The present study shows a more prominent antigen presentation via MHC class I and upregulation of endosomal/vacuolar pathway important for loading antigenic peptides to MHC class I. Antigen presentation via MHC class II was not significantly represented by the upregulated proteins. This is contrary to the previously held belief that the presence of MHC class II and not class I is important for resistance to leishmaniasis [21]. The antigens presented may be originating from the *Leishmania* species itself or an endobiont virus or both. This study suggests that exogenous antigens derived from *L. donovani* may be presented via pathways dependent and independent of TAP [22].

Moreover, the significant up-regulation of Calnexin in this study suggests a potential involvement of phagosome biogenesis in the host-pathogen interactions of CL in Sri Lanka. Phagosomes are intracellular membrane-bound compartments linked to the endoplasmic reticulum, providing a protective environment where *L. donovani* can evade acquiring lysosomal properties [23].

Gene expression studies done on the cytokine response by our group [24] and other groups [25] have revealed the presence of an up-regulated Th1 response in CL in Sri Lanka. The current study on proteome profiling of CL lesions has further proven the presence of an up-regulated Th1 response in *L. donovani* induced CL in Sri Lanka at the post translational stage. Four proteins associated with the IFN-γ signaling pathway were seen to be significantly upregulated in patient tissues compared to healthy controls and six proteins in the IL-12 signaling pathway were also significantly upregulated. The action of IL-12 is important in bridging the innate and adaptive arms of the host immune response [26]. The IL-12 family of cytokines consists of IL-12 and IL-23 which are pro-inflammatory, and IL-27 and IL-35 which are immune-suppressive and regulate the immune response in autoimmune and infectious diseases [27]. Pathway analysis in this study has shown the presence of IL-27 and Il-35 signaling pathways, which need to be validated and quantified further to decipher their extent of involvement in the immune response against CL.

In addition, processes involved in the protein translation in ribosomes and cytoplasmic transport are also significantly up-regulated indicating the active inflammatory milieu created in the lesion compared to the healthy skin. Also, the current study reveals a notable up-regulation in the pathway of apoptosis-induced DNA fragmentation. Even though histological evidence of apoptosis or necrosis is not very prominent in CL due to *L. donovani* in Sri Lanka [28], this study points to the presence of apoptosis, possibly at a low magnitude in the lesions.

### Conclusion

*Leishmania spp*. parasites comprise a diverse group of protozoans that lead to a range of disease manifestations. The immunological responses to these parasites are highly intricate. This complexity hinges on factors like the causative species and possibly the strain involved. In Sri Lanka, CL has emerged as an established vector-borne parasitic disease, characterized by a fascinating clinical presentation. This study which investigates the proteomic profiling of leishmaniasis lesions revealed a multitude of probable immunological and pathological mechanisms operating in patients with CL in Sri Lanka such as the unfolded protein response, a probable association of an endosymbiont virus in the parasite, IFN-α/β signaling, and phagosome biogenesis, which need to be further elaborated using more in-depth and targeted investigations.

## Methods

### Sample collection and confirmation of diagnosis

This study received ethical approval from the Ethics Review Committee of the Faculty of Medicine, University of Kelaniya, Sri Lanka (P/99/06/2013) and was conducted adhering to the approved protocol and in agreement with the Helsinki Declaration. Patients and controls were recruited voluntarily and informed written consent was obtained before sample collection.

Patients with skin lesions suspected of CL were recruited from Base Hospital Padaviya and the Sri Lanka Army. Lesion biopsies with a diameter of 3-4 mm were obtained from the active edge of the CL lesion before starting treatment. The diagnosis was established with light microscopy of Giemsa-stained tissue impression smears and species diagnosis was confirmed using previously established molecular methods [29]. Control skin specimens were obtained from incision sites of patients with no signs or symptoms of leishmaniasis, who underwent minor surgical procedures due to unrelated surgical causes. Skin biopsy specimens were immediately submerged in RNAlater and stored at −20 ^0^C until further analysis. Eight patient and eight control skin specimens were processed for proteomic profiling by mass spectrometry.

Samples for immunohistochemical (IHC) validation of the unfolded protein response (UPR) pathway were selected from previously archived formalin fixed paraffin embedded lesion and control specimens. Thirty lesion specimens from leishmaniasis-confirmed patients and six control specimens from patients undergoing minor surgical procedures for unrelated surgical causes were used for IHC staining. This part of the study received ethical approval from the Ethics Review Committee of the Faculty of Medicine, University of Kelaniya, Sri Lanka (Ref. No. P/21/03/2021)

### Sample processing for proteomics

Sample preparation was carried out at the Campus Chemical Instrument Center (CCIC) Mass Spectrometry and Proteomics Facility, Ohio State University, Columbus, Ohio, USA. Sample preparation was done in a Class II type A2 biosafety cabinet (NuAir, Minnesota, USA). Tissue samples stored in RNAlater were removed from the reagent, blotted on a filter paper to remove traces of RNAlater, and placed in new 1.5 ml microcentrifuge tubes (Fisher Scientific, New Hampshire, USA). To each of the tubes, 100 uL of 0.2 % RapiGest SF Protein Digestion Surfactant (Waters, Milford, MA, USA) in 50 mM NH_4_HCO_3_ was added. Samples were sonicated in a Sonic Dismembrator (Fisher Scientific, New Hampshire, USA) at speed 6 for 4 times and speed 5 twice, each sonication lasting 3 seconds. Sonicated samples were then heated at 105 °C in a heat block for 30 min, following which they were cooled on ice for 5 minutes. Samples were then vortexed for 5 minutes, following which they were heated at 70 °C for 2 hours.

Dithiothreitol (ThermoFisher Scientific, USA) was added at a final concentration of 5 mM to reduce the disulphide groups and maintain sulfhydryl (-SH) groups which would make protein fragmentation and analysis more effective. Samples were then heated at 60 °C for 30 minutes, following which, iodoacetamide (Acros Organics, NJ, USA) was added at 15 mM final concentration and incubated at room temperature, in the dark for 15 minutes to inhibit proteases. For digestion of proteins, 1 µg of sequencing grade trypsin (Promega, Wisconsin, USA) was added and samples were incubated at 37 °C overnight.

Rapigest was precipitated by adding trifluoroacetic acid (Fisher Scientific, New Hampshire, USA) to a final concentration of 0.5% and incubating at 37 °C for 30 minutes. Samples were then centrifuged at 13,000g for 15 minutes. The supernatant was then transferred to a microcentrifuge tube (Eppendorf®) and dried in a speed vac (Eppendorf Vacufuge Plus, Hamburg, Germany). Samples were stored at −80 °C until analysis. They were then re-suspended in 50 mM acetic acid (Ultrex II Ultrapure Reagent, J.T. Baker ™) and peptide concentrations were determined from their absorbance at 280 nm using a Nanodrop 1000 spectrophotometer (Thermo Fisher Scientific, USA).

### Instrument protocol-tandem mass spectrometry

Prior to tandem mass spectrometry (MS^2^), samples were subjected to two –dimensional liquid chromatography (2-D LC) separation using a Thermo Scientific 2D rapid separation liquid chromatography (RSLC) high-pressure liquid chromatography (HPLC) system. A sample volume consisting of 12 ug of peptides was first separated on a 5 mm x 300 μm Ethylene Bridged Hybrid (BEH) C_18_ column with 5 μm particle size and 130 Å pore size. Solvent A was composed of 20 mM ammonium formate (Fisher Scientific New Hampshire, USA) at pH 10, and solvent B was 100% HPLC grade acetonitrile (Sigma Aldrich, Missouri, USA). Peptides were eluted from the column in eight successive fractions using 9.5, 12.4, 14.3, 16.0, 17.8, 19.7, 22.6 and 50% solvent B. Each eluted fraction was then trapped, diluted, neutralized, and desalted on a µ-Precolumn Cartridge (Thermo Fisher Scientific) for the second-dimension separations performed with a 15 cm x 75 cm PepMap C18 column (ThermoFisher Scientific, Waltham, MA) with 3 μm particle size and 100 Å pore size. For the Thermo Scientific 2D RSLC HPLC system, the flow rate for the analytical column was 500 μL/min. The gradient was 0 to 5 min, 2% solvent B; 5 to 38 min, 35% solvent B; 38 to 46 min, 35-55% solvent B; 46 to 47 min, 55-90% solvent B. Mobile Phase B was kept at 90% for 1 min before quickly brought back to 2%. The system was equilibrated for 11 min for the next separation.

Tandem mass spectrometry data was acquired with a spray voltage of 1.7 KV and the capillary temperature used was 275 °C. The scan sequence of the mass spectrometer was based on the preview mode data dependent TopSpeed™ method: the analysis was programmed for a full scan recorded between *m/z* 400 – 1600 and an MS^2^ scan to generate product ion spectra to determine amino acid sequence in consecutive scans starting from the most abundant peaks in the spectrum in the next 3 seconds. To achieve high mass accuracy mass spectrometry determination, the full scan was performed in Fourier Transformation (FT) mode and the resolution was set at 120,000. The automatic gain control (AGC) target ion number for the FT full scan was set at 2 × 10^5^ ions, the maximum ion injection time was set at 50 ms, and micro scan number was set at 1. Tandem mass spectrometry was performed using ion trap mode to ensure the highest signal intensity of MS^2^ spectra using both collision-induced dissociation (CID) for 2+ and 3+ charges and electron-transfer dissociation (ETD) for 4+ to 6+ charges. The AGC target ion number for the ion trap MS^2^ scan was set at 1000 ions, the maximum ion injection time was set at 100 ms, and micro scan number was set at 1. The CID fragmentation energy was set to 35%. Dynamic exclusion is enabled with an exclusion duration of 15 with a repeat count of 2 within 30s and a low mass width and high mass width of 10 ppm.

Sequence information from the MS^2^ data was processed by converting .raw files into a mgf file using MS convert (ProteoWizard) and then mgf files from each of the fractions was merged into a merged file (.mgf) using Merge mgf (ProteinMetrics). Isotope distributions for the precursor ions of the MS^2^ spectra were de-convoluted to obtain the charge states and mono-isotopic *m/z* values of the precursor ions during the data conversion. The resulting mgf files were searched using Mascot Daemon by Matrix Science version 2.5.1 (Boston, MA, USA) and the database was searched against the human database. The mass accuracy of the precursor ions was set to 10 ppm, the accidental pick of one ^13^C peak was also included in the search. The fragment mass tolerance was set to 0.5 Da. Considered variable modifications were oxidation (Methionine), deamidation (Asparagine and Glutamine), acetylation (Lysine), and carbamidomethylation (Cysteine) was considered as a fixed modification. Four missed cleavages for the enzyme were permitted. A decoy database was also searched to determine the false discovery rate (FDR) and peptides were filtered according to the FDR. Only proteins identified with <1% FDR as well as a minimum of 2 peptides were reported.

### Bio-informatics analysis of proteomics data

For this analysis raw data on MS/MS spectral counts were used. If a protein had a spectral count of < 6 in ≥ 90% of samples that protein was filtered out from the data analysis [30]. After filtering, 388 protein identities were left for further analysis. The Voom normalization was applied to normalize the data across all samples to reduce the bias in signal intensities from run to run. Comparison between groups was done by the ‘analysis of variance’ (ANOVA) method. The p-value obtained was adjusted for multiple corrections using the Benjamini-Hochberg procedure. All the proteins with an adjusted p-value <0.01 were considered as significantly expressed between the groups compared. Significantly expressed proteins thus identified were entered into the UniProt human database [31] and converted to their corresponding gene names. Protein-protein interactions were assessed using the database ‘STRING: functional protein association networks’, Version 10.5 (https://string-db.org/) [32]. Pathway analysis was done using the Reactome pathway portal, version 3.2 (http://www.reactome.org) [33].

### Immunohistochemical validation of IRE1, ATF6 and PERK

Immunohistochemical staining for IRE1, ATF6, and PERK were performed on paraffin-embedded tissue samples in all selected cases and controls. Tissue sections of 4 µm were cut using a microtome and the sections were mounted on positively charged slides and dried overnight in an oven at 60^0^ C. The slides were dewaxed in xylene and rehydrated with 100% ethanol and 90% ethanol for 10 minutes each. The slides were then washed with deionized water two times for 5 minutes. The slides were subjected to a 25-minute microwave-boiling process for antigen retrieval, using an appropriate buffer and pH as specified in Table 3.

**Table 3:** Immunohistochemical protocols and scoring for IRE1, PERK, and ATF6 markers.

| Marker | Supplier & Clone | Antigen retrieval | Primary dilution | Positive control | Interpretation | Scoring |
| --- | --- | --- | --- | --- | --- | --- |
| IRE1 | Abcam ab37073 | Microwave in pH 6.0 citrate buffer | 1:500 | Small intestine | Cytoplasmic staining | Intensity score* of cytoplasmic staining × Proportion score** of cytoplasmic staining |
| PERK | Abcam ab79483 | Microwave in pH 9.0 EDTA buffer | 1:40 | Colonic carcinoma | Cytoplasmic staining and nuclear staining | (Intensity score of cytoplasmic staining × Proportion score of cytoplasmic staining) + (Intensity score of nuclear staining × Proportion score of nuclear staining) |
| ATF6 | Abcam ab203119 | Microwave in pH 6.0 citrate buffer | 1:250 | Kidney | Nucleus and cytoplasmic staining | (Intensity score of cytoplasmic staining × Proportion score of cytoplasmic staining) + (Intensity score of nuclear staining × Proportion score of nuclear staining) |

\*Intensity score = 0 - no staining; 1-weak staining; 2- moderate staining; 3-strong staining
\*\*Proportion score = 0 - 0-5% cells; 1 - 6-30% cells; 2, - 31-70% cells, 3- 71-100% cells

Endogenous peroxidase was blocked by incubation in 3% hydrogen peroxide for 10 min followed by washing in deionized water and wash buffer (1× TBST). Non-specific binding was blocked by adding the blocking solution for 1 hour at room temperature in a humidified chamber. The tissue sections were then incubated overnight at 4^0^ C with primary antibodies.

The antigen-antibody complex was detected by the Labeled Streptavidin–Biotin (LSAB) staining method using a biotinylated goat anti-rabbit antibody (DakoCytomation), subsequently conjugated with streptavidin-horseradish peroxidase (HRP) and visualized by reacting with 3,3′-diaminobenzidine for color detection. The tissue sections were counterstained with hematoxylin. Dehydration was done by submerging the slides in 95% ethanol, 100% ethanol, and xylene twice for 10 min each respectively. Finally, the sections were mounted using a mounting medium and observed under the microscope.

The IHC-stained tissue samples were evaluated by light microscopic examination for the expression of IRE-1, PERK and ATF-6. The intensity of staining for each of the markers and the proportion of cells that expressed attaining were evaluated in 5 random fields (400× magnification) for each section. IRE-1 was assessed for cytoplasmic staining. PERK and ATF-6 were evaluated for both cytoplasmic and nuclear staining. The proportion score was calculated as: 0, 0-5% cells were stained; 1, 6-30% cells were stained; 2, 31-70% cells were stained, and 3, 71-100% cells were stained. The staining intensity was scored as follows: 0, no staining; 1 weak staining; 2, moderate staining; 3, strong staining [34].

The overall score for IRE-1 was calculated as:

Intensity score for cytoplasmic staining × Proportion score cytoplasmic staining The overall score for ATF-6 and PERK was calculated as [34,35]:

(Intensity Score for nucleus staining × proportion score for nucleus staining) + (Intensity score for cytoplasmic staining × Proportion score cytoplasmic staining)

The cases that showed staining for each of the markers, IRE1, PERK, and ATF-6 were further categorized as low positivity and high positivity based on the score obtained for each of the markers. IRE −1 – low positive <= 4 and high positive > 5, PERK – low positivity <= 8 and high positivity >9, ATF-6 – low positivity <= 2 and high positivity >3. The selection of these arbitrary cut-off values is made specifically for this study, as there is no uniform threshold established in the relevant literature.

### Statistical analysis

Statistical analyses were carried out using SPSS (version 25.0, SPSS Inc, Chicago, IL. USA) software. The association between the degree of staining for each marker and the clinical and pathological features was assessed using the Chi-square test. A *P* value < 0.05 was considered statistically significant.

## Acknowledgments

This work was supported by the National Institute of Allergy and Infectious Diseases of the National Institutes of Health [grant numbers R01AI099602 and UO1A136033 to N.D.K.]; the University Grants Commission of Sri Lanka [grant number UGC/VC/DRIC/PG/2013/KLN/03]; and the University of Kelaniya, Sri Lanka [grant number RP/03/04/06/01/2014]. The content is solely the responsibility of the authors and does not necessarily represent the official views of the National Institutes of Health or any other funding agency.

The authors would like to acknowledge Dr. B.N.S. Jayawardhana and N. L. Munasinghe for assistance with the collection of control specimens; Prof. Abhay Satoskar and Dr. Steve Oghumu for guidance in mass spectrometry-based proteomic work; Mrs. Sandya Liyanarachchi of Ohio State University for helping with the bioinformatic analysis of proteomics data and all the patients for participating.

## Supporting information

**S1 Table.** Enriched biological processes (GO). **S2 Table.** Enriched molecular functions (GO). **S3 Table.** Enriched cellular component (GO).

**S4 Table.** Kyoto Encyclopedia of Genes and Genomes (KEGG) pathways.

**S5 Table.** Pathways showing a significant association with the upregulated proteins as determined by the ‘Reactome Pathway Portal’, version 3.2.

